# X-Ray contrast-adjustable 3D printing for multimodal fusion of microCT and histology

**DOI:** 10.1101/2025.11.21.689680

**Authors:** Philipp Nolte, Chris Johann Ackurat, Marcel Brettmacher, Marius Reichardt, Marieke Stammes, Christoph Rußmann, Christian Dullin

## Abstract

**Object:** Phantoms and reference structures are essential tools for calibration and correlative imaging in pre-clinical and research applications of X-Ray-based imaging. They serve as reference standards, ensuring consistency and accuracy in imaging results. However, generating individual phantoms often involves a complex creation process, high production costs, and significant time investment.

**Method:** Conic reference structures were 3D printed using a mixture of UV-curable resin and X-Ray contrast agents. These structures were then embedded together with lung specimens of SARS-CoV-2-infected rhesus macaques in a methyl methacrylate-based solution. The polymerized blocks were scanned using propagation-based phase-contrast microCT, a method chosen for its superior ability to enhance contrast especially in low-absorbing biological samples. Utilizing the conic reference structures, subsequently performed histological sections were co-registered into the 3D context of the microCT data sets.

**Result:** The produced 3D-printing models were highly visible in terms of contrast and detail in both imaging methods, allowing for a precise co-registration of microCT and histological imaging.

**Conclusion:** The novel methodology of using contrast agents and resin in 3D-printing enables the generation of customizable, contrast-specific phantoms and reference structures. These can be straightforwardly segmented from the embedding material, significantly simplifying and enhancing the workflow of multimodal imaging processes. In this study, 3D printed conic reference structures were effectively used to automate and streamline the precise multimodal fusion of microCT and histological imaging.

## 1 Introduction

Correlative imaging has emerged as a prominent research focus in the analysis of biomedical tissue samples, primarily due to its ability to fuse complementary datasets that were traditionally assessed separately, thereby enhancing diagnostic accuracy. Typically, tissues are processed for microscopic examination via standard histological workflows, which involve embedding the specimen in a supportive matrix, sectioning it into thin slices, staining, and imaging under a microscope. The resulting two-dimensional (2D) whole slide images (WSIs) offer high specificity but lack contextual three-dimensional (3D) structural information. To address this limitation, researchers have developed approaches to align WSIs with micro-computed tomography (microCT) scans acquired prior to sectioning, thereby extending histological analysis into the third dimension [1–4]. This alignment is commonly achieved through image registration, which generally requires intensive computational power due to the difficulty of accurately locating the 2D histological slice within the 3D volume. Albers et al. [1] proposed a manual method based on visual similarity for coarse alignment. In contrast, Chen et al. [2] developed a deep learning-based approach to automate this task.

Recently, 3D printing has been introduced as a versatile tool to assist the histological routine, for example by the creation of sectioning guides [5–8] or for the printing of anatomical models [9]. In our previous work [10], we presented the use of extrinsic 3D printed conical markers to aid in aligning WSIs with microCT scans of hard tissue specimens embedded in resin. These markers were fabricated using a binder-jetting process with a calcium sulfate powder, resulting in high-density structures with coarse texture that were clearly visible in both microCT and histological images. However, the markers were characterized by low printing resolution and exhibited frayed edges upon sectioning. Alternative 3D-printing methods such as fused deposition modeling (FDM) and digital light processing (DLP), as well as their typical materials, were found unsuitable due to their resulting X-Ray contrast being similar to or lower than the embedding resin, rendering them invisible in the scan.

To enhance the resolution and precision of our conic marker-based workflow, we developed a novel approach combining UV-curable resin, commonly used in DLP 3D-printing, with iodine-based X-Ray contrast agents. This methodology enables the fabrication of high-fidelity phantoms or fiducial markers with tunable radiopacity, thereby allowing accurate co-registration between histological images and highresolution X-Ray imaging modalities. Leveraging the superior resolution of DLP printing, we produced finely detailed markers that are both structurally robust and radiographically visible. This improvement allowed to fuse high-resolution scans obtained from synchrotron-based propagation-based phase-contrast micro-computed tomography (PBI-microCT) with corresponding histological sections. The result was a reduction in the difference in spatial resolution between histology and PBI-microCT by a factor of 20, enabling the generation of multimodal datasets of the specimen, while achieving high similarity.

The samples containing both the tissue specimens and the conic reference markers were processed using a combination of automated sectioning and laser microtomy, as described in our previous work [11]. This method enabled fast and precise sectioning of resin-embedded specimens. When Integrated with our proposed 3D printing approach, this allowed the automation of the entire preparation and multimodal analysis pipeline for resin-embedded samples. Segmentation of the conic reference structures in both imaging modalities was performed using (i) traditional image processing techniques and (ii) the Segment Anything Model (SAM) [12, 13]. While the latter significantly streamlined the segmentation process and improved robustness, it requires GPU resources for efficient execution. Additionally, we generated comprehensive multimodal correlative datasets, which have potential utility for training machine learning models in the domain of digital pathology, particularly for tasks involving multimodal image registration, tissue classification, or structure identification.

Here we present a new 3D printing method for the generation of models with specified X-Ray contrast. The proposed method is used to create conic-markers, which allow for the precise, rapid and simplified alignment of both PBI-microCT scans and WSI.

## 2 Methods

### Design and fabrication of the 3D-printed reference markers

The reference markers were designed as conical structures to enable precise localization of 2D histological sections within the corresponding 3D microCT datasets, as previously demonstrated in [10]. To improve upon this design, three cones were connected to a common base, simplifying the embedding process and enhancing positional stability. Each cone was designed in CAD software (Creo, PTC Inc.) with a height of 11 mm, a base diameter of 3 mm, and an apex diameter of 0.5 mm. The corresponding STL file can be accessed at https://github.com/devphilno/3DPrint-CT-Histo. In our earlier work [10], the reference cones were fabricated from calcium sulfate powder using a binder-jetting process. Although functional, this method produced coarse surface textures and frayed edges during sectioning. In the present study, DLP 3D printing (Phantom Mono 4K, Shenzhen Anycubic Technology Co., Ltd.) was used to achieve the high geometric fidelity required to match the resolution of PBI-microCT imaging. To improve radiopacity and thus enable separation from the embedding medium during segmentation, the UV-curable resin (Water-Washable Resin, ELEGOO EU) was mixed with an iodine-based X-ray contrast agent (Ultravist 370, Bayer Vital GmbH) prior to printing. This combination ensured both structural precision and high X-ray visibility.

### Specimen preparation and embedding protocol

Biopsy punches (8 mm in diameter) from formalin-fixed rhesus macaques (Macaca mulatta) lungs were stained with phosphotungstic acid as described by Saccomano et al. [14] and embedded, together with cone-shaped reference phantoms, in a methyl methacrylate (MMA)-based resin (Technovit 9100, Kulzer GmbH) using 25 mm-wide molds. Embedding was performed following the manufacturer’s protocol

### MicroCT acquisition

After polymerization, the specimens were imaged at the Synchrotron Radiation for Medical Physics (SYRMEP) beamline of the Elettra synchrotron facility in Trieste, Italy [15]. Imaging was conducted using a white-beam configuration in propagation-based phase-contrast imaging (PBI) mode, with a sample-to-detector distance of 150 mm. The reconstructed pixel size was 3.95 *µ*m, yielding an effective field of view of approximately 7 *×* 4 mm^2^. Prior to reconstruction using the filtered back-projection algorithm, Paganin’s phase retrieval algorithm [16] was applied with a delta-to-beta ratio of 50, both implemented in the SYRMEP Tomo Project (STP) software [17]. The resulting 3D datasets were then stitched using NR-Stitcher [18] to obtain continuous volumetric reconstructions. Image type conversion and contrast range adjustments were performed using Fiji [19].A subset of the blocks were also scanned using an *in vivo* microCT system (Quantum GX, Revvity Inc.) with the following settings: tube voltage of 90 kV, tube current of 88 *µ*A, field of view (FOV) of 36 *×* 36 mm^2^, and a total acquisition time of 4 min, resulting in an isotropic resolution of 36 *µ*m. The region of interest was subsequently upscaled using the microCT viewer, achieving a FOV of 18.432 *×* 18.432 mm^2^ and a voxel size of 18 *µ*m.

### Laser microtomy, histological staining and microscopy

A priming cut to expose the target histological region was performed using a pathological saw (Cut-Grinder Primus, Walter Messner GmbH). The trimmed blocks were then mounted to microscope glass slides (X-tra-Adhesive, Leica Biosystems Nussloch GmbH) and sectioned using a laser microtome (Tissue Surgeon, LLS Rowiak LaserLabSolutions GmbH). The resulting histological sections were stained according to the Sirius Red protocol [20, 21] and subsequently imaged with a microscope (LSM 700, Carl Zeiss Microscopy GmbH) resulting in a pixel size of 1.16 µm.

### Software development

The majority of the software was developed in a remote environment and subsequently executed on a GPU server (HGX A100, NVIDIA Corp.). Image processing and analysis were carried out using Python 3.10.12, with the following libraries: numpy 1.24.4 [22], pandas 2.2.2 [23], scikit-image 0.25.2 [24], Pillow 10.4.0 [25], and OpenCV 4.7.0 [26]. AI-based segmentation was performed using SAM 2 [13]. Image registration was implemented using simpleITK 2.5.0 [27, 28]. All Python development was conducted in Jupyter Notebooks [29, 30]. The code is available at https://github.com/devphilno/3DPrint-CT-Histo. The calculation of the similarity scores was performed using the LitSHi registration tool [31].

## 3 Results

The integration of the 3D-printed conic reference enabled successful fusion of 3D microCT and 2D histology. Since the resulting X-Ray absorption of the phantoms was calibrated specifically during printing by using defined mixes of resin and iodine based X-Ray contrast agents, identification of the specimen and reference structure was simplified.

### General workflow

Reference structures were fabricated using a DLP printing process, employing a mixture of UV-curable resin and iodine-based contrast agent and (Fig. 1A). Both the lung tissue specimen and the 3D-printed reference structures were embedded in resin (Fig. 1B). Due to the increased absorption resulting from the iodine-based contrast agent in both the tissue and the 3D-printed parts, the image contrast in phase-contrast X-Ray PBI-microCT (Fig. 1C) was enhanced, thereby enabling straightforward segmentation by applying adaptive thresholding. Following scanning, the resin blocks were sectioned, and histological staining was performed (Fig. 1D). Each resulting histological slide was registered within the 3D microCT volume by analyzing the sections of the conic reference structures, as described in our earlier work [10]. The associated microCT plane was then extracted and fused with the histological section (Fig. 1E). The complete workflow is illustrated in Fig. 1.

**Fig. 1.**
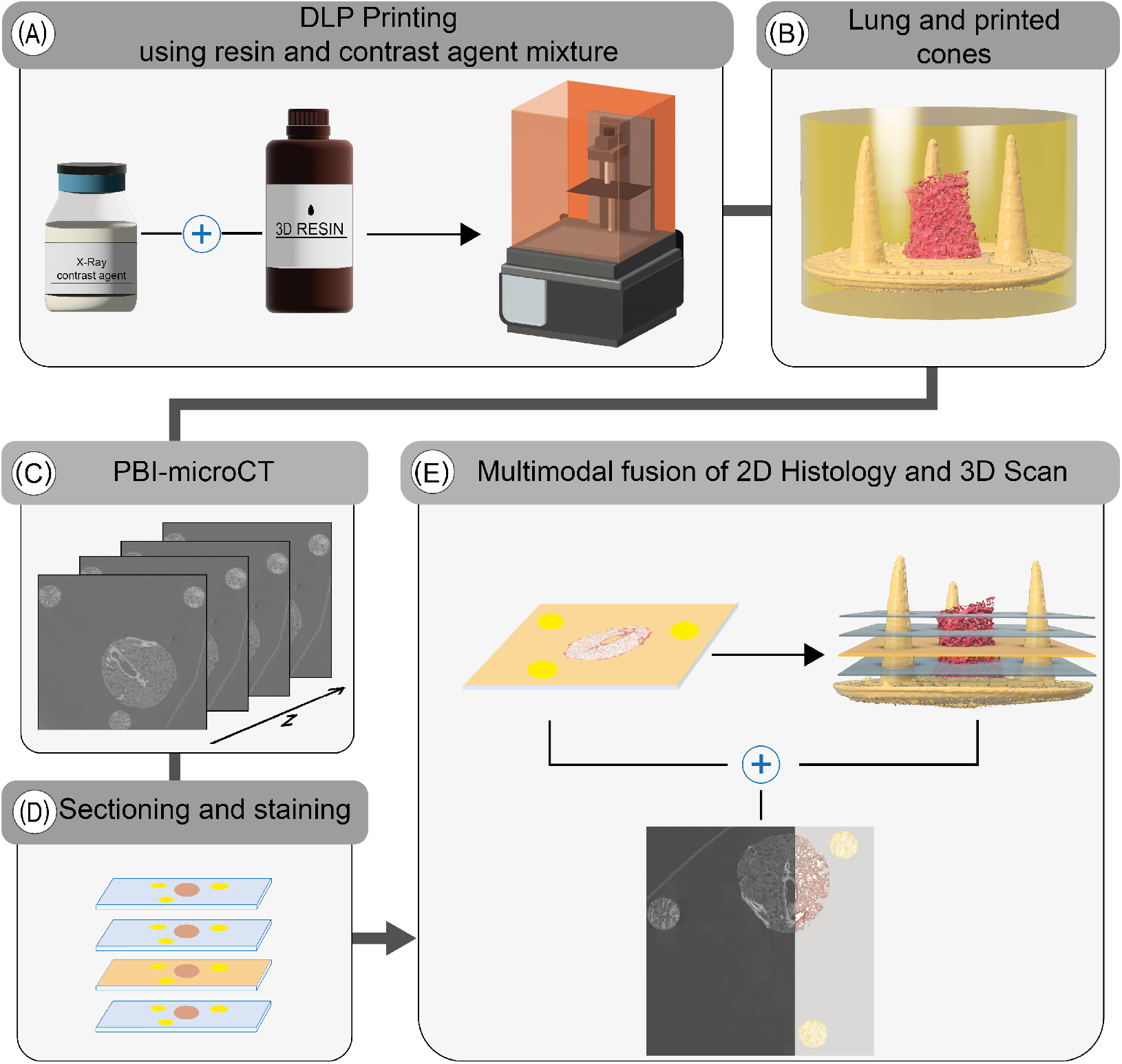
Overview of the extended histological routine. Reference structures are first printed using a tunable UV-resin mixed with a contrast agent (A). The resulting conical markers are embedded alongside the tissue specimen (lung punch biopsy, in this case) in resin (B). The polymerized block is then scanned using PBI-microCT (C) and subsequently sectioned (D). After histological staining, the position of each individual 2D slice is determined using the conical markers and precisely registered to the corresponding *in silico* plane of the 3D volume (E), thereby facilitating a multimodal fusion of both datasets.

### Crafting of the resin-contrast agent mixture and DLP printing

In order to enhance X-Ray absorption and achieve high contrast relative to the embedding medium (MMA), standard UV-curable resin was mixed with an iodine-based contrast agent commonly used to improve the visibility of non-mineralized soft tissue in both human and animal imaging [32, 33]. DLP printing was employed to overcome the limited spatial resolution and coarse structural fidelity observed in previous reference structures produced via binder jetting with calcium sulfate, as described before [10]. Although DLP printing was available as a higher-resolution alternative for our previous work, the resulting polymerized resin structures exhibited low contrast relative to the embedding resin, rendering them visually indistinguishable. By incorporating the iodine-based contrast agent into the printing resin, it became possible to tune the resin’s X-Ray attenuation properties, thereby enabling clear segmentation from the surrounding medium. For the absorption-based imaging the X-Ray contrast is commonly quantified using CT numbers, expressed in Hounsfield Units (HU), which assign a standardized scale to pixel intensities. This scale ranges approximately from -1000 HU for air to over 500 HU for bone, and can exceed 3000 HU for metals [34, 35]. While HU values are approximated in standard microCT systems due to the use of polychromatic X-Ray sources [36, 37], they still provide a practical basis for material differentiation. To estimate the required amount of contrast agent, we used the following equation:

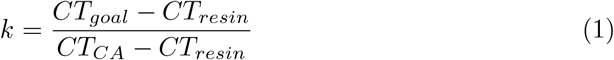

Here, *k* represents the mixing coefficient, *CT*_*goal*_ is the target CT number, *CT*_*resin*_ is the attenuation of the pure resin, and *CT*_*CA*_ is the attenuation of the pure contrast agent. The achievable range of attenuation is constrained by the intrinsic contrast values of both the resin and the contrast agent.For the creation and validation of the phantoms, a preliminary scan using a classical absorption-based microCT was used (see Fig. 2). While the resulting images are characterized by significantly lower spatial resolution, it still allowed for a quality check of the overall contrast agent distribution inside the reference structure. In our experiments, we obtained CT numbers ranging from about 180 HU to 4600 HU. The mixture was composed of 18% contrast agent. Two representative slices from the absorption-based microCT scan are shown in Fig. 2. Fig. 2A presents a vertical slice containing both the cone and lung tissue. Variations in contrast are visible within the 3D-printed structure (indicated by the white arrows), which can be attributed to sedimentation of the contrast agent during the DLP printing process. For comparison, Fig. 2B shows a horizontal slice, providing a visual reference for correlation with the PBI-microCT images processed in this study. Notably, no modifications to the DLP printer’s default settings were required, making this method easily integrable into existing printing workflows.

**Fig. 2.**
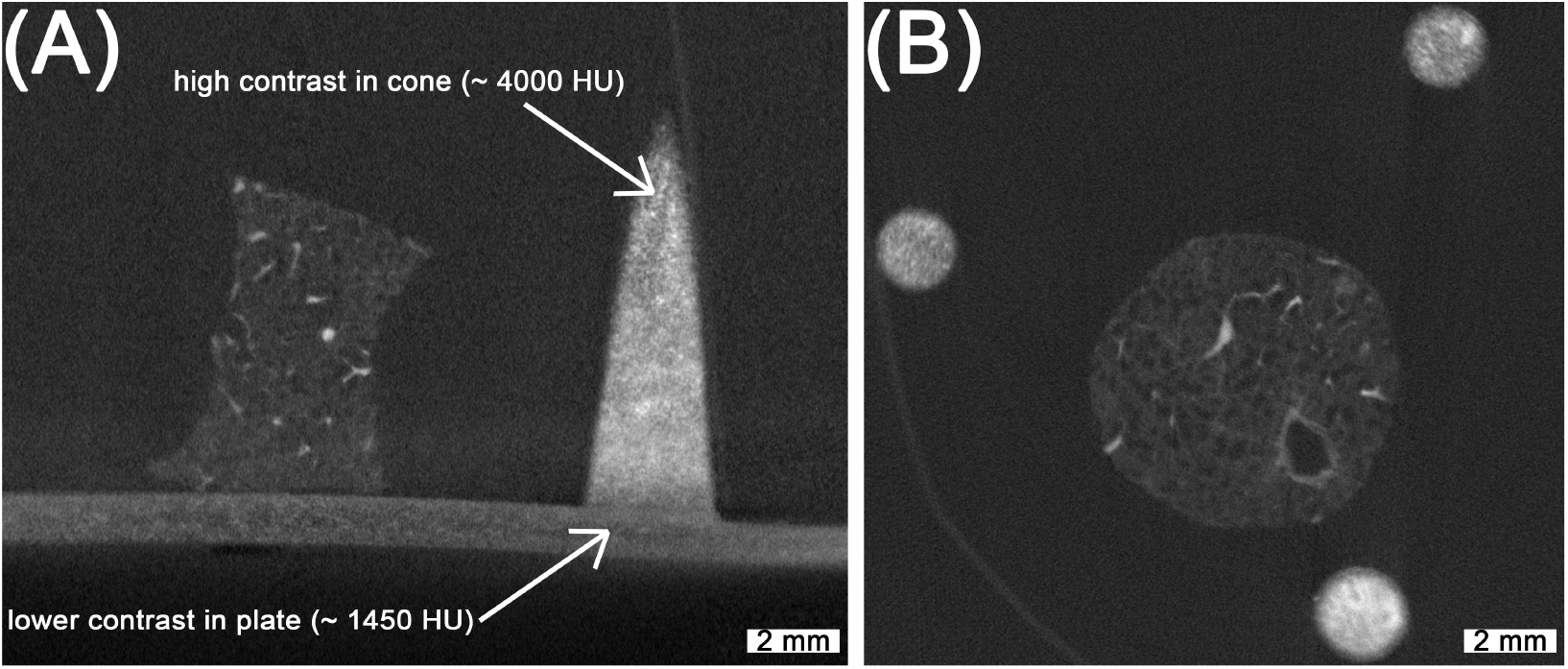
Absorption-based microCT scan of the lung with 3D-printed reference structures. While the overall high contrast achieved allows for straightforward segmentation, non-uniform distribution of the contrast agent is evident (as indicated with the white arrows). The spatial resolution in these scans is noticeably lower, as illustrated in (B), complicating the precise overlay of fine branches in the lung tissue.

### Design of the PBI-microCT analysis pipeline

After scanning, individual image stacks corresponding to each field of view were reconstructed and stitched together. The pixel value range was adjusted based on the global minimum and maximum values of the microCT stack before stitching. Following this preprocessing step, a subset of 500 sampled images was selected from the full stack of 1982 layers for analysis. This layer-by-layer approach deviated from the previously introduced methodology, where the axis of inertia for each cone was detected directly in the 3D volume. Given the significantly larger dataset in this study, this sampling strategy was adopted to improve computational efficiency. As illustrated in Fig. 3A the selected images were processed in a multi-step, rule-based procedure: Adaptive Thresholding [38], Edge Filtering based on a Canny edge detection [39], morphological opening and closing, Filtering and Ellipse fitting to i) detect the conic sections resulting from the cut conic reference structures and ii) extract the height and width as well as the x- and y-coordinates of the center point of the fitted ellipse. Due to imaging artifacts, the detection process produced false positives, as shown in Fig. 3B), where sections where associated with the wrong cone. These outliers were removed based on their distance from the mean of the clusters and the detected width of the ellipse as shown in Fig. 3C.

**Fig. 3.**
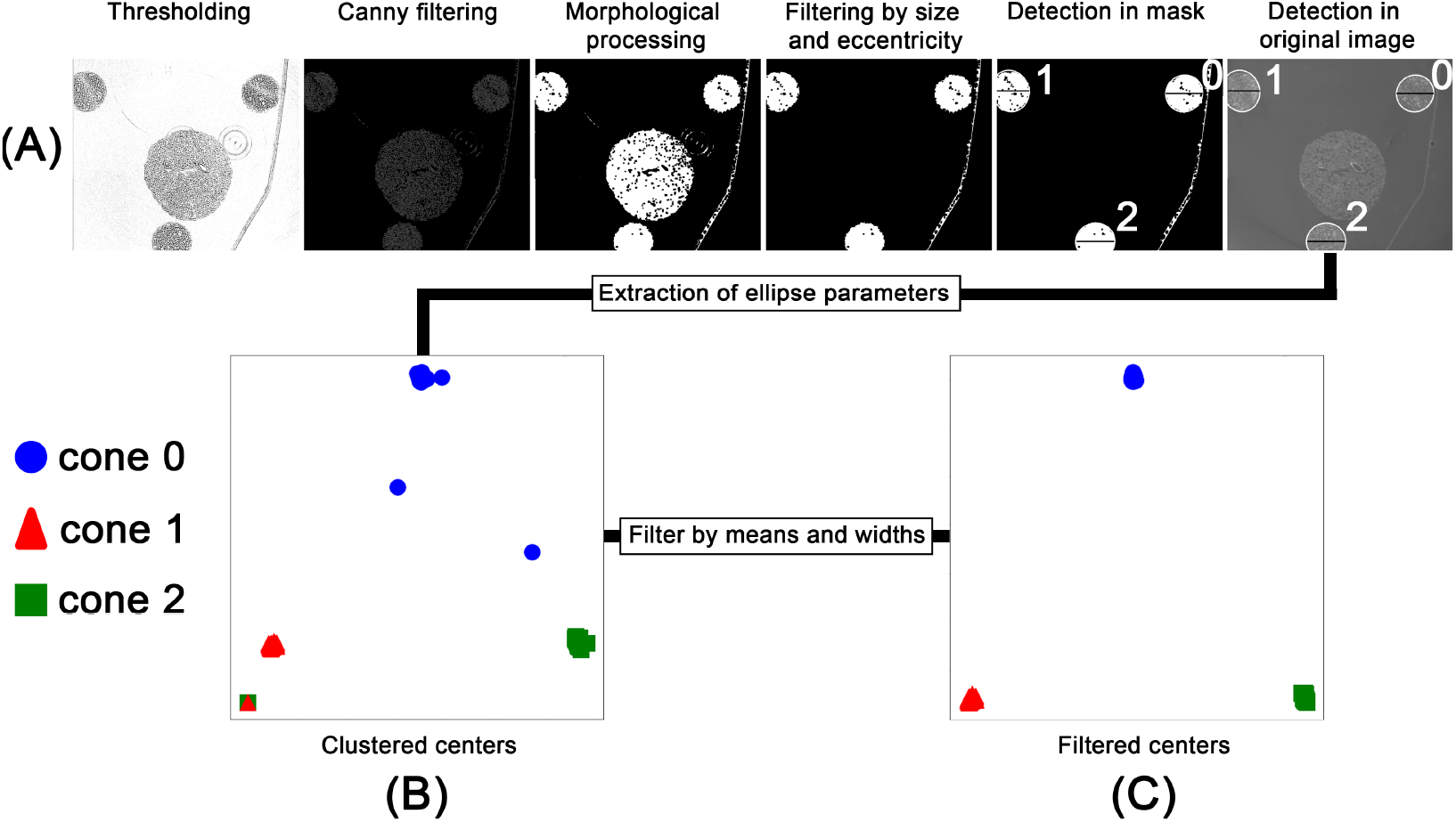
Workflow for the rule-based processing of the PBI-microCT images. After thresholding and edge detection, the images were morphologically processed to generate an object mask. This mask served as input for an ellipse-fitting algorithm. The fitted ellipses were filtered based on size and eccentricity and overlaid on the original image (A). The extracted ellipse parameters, namely, center coordinates and width, were saved and clustered (B). These clusters were then filtered based on their density and ellipse width (C).

The rule-based detection of the cones and extraction of ellipse parameters resulted in a detection rate of approximately 75% for the 500 images processed. Although missing data points could be interpolated, the resulting relationship between cone width and height would reduce precision. Therefore, we employed the SAM2 model [13] to detect the cones in each image. As SAM provides zero-shot generalization, multiple objects, such as tissue regions and artifacts, were also detected. However, using the mean center coordinates obtained from the filtered rule-based detection step, SAM detections corresponding to the conic sections were segmented (see Fig. 4A). The resulting binary mask depicted in Fig. 4B allowed for severely simplified detection of the individual cone and subsequent computation of the width of each cone. For each cone, the inverse relationship between width and height was mapped and smoothed using a regression function. The resulting graphs are shown in Fig. 4C.

**Fig. 4.**
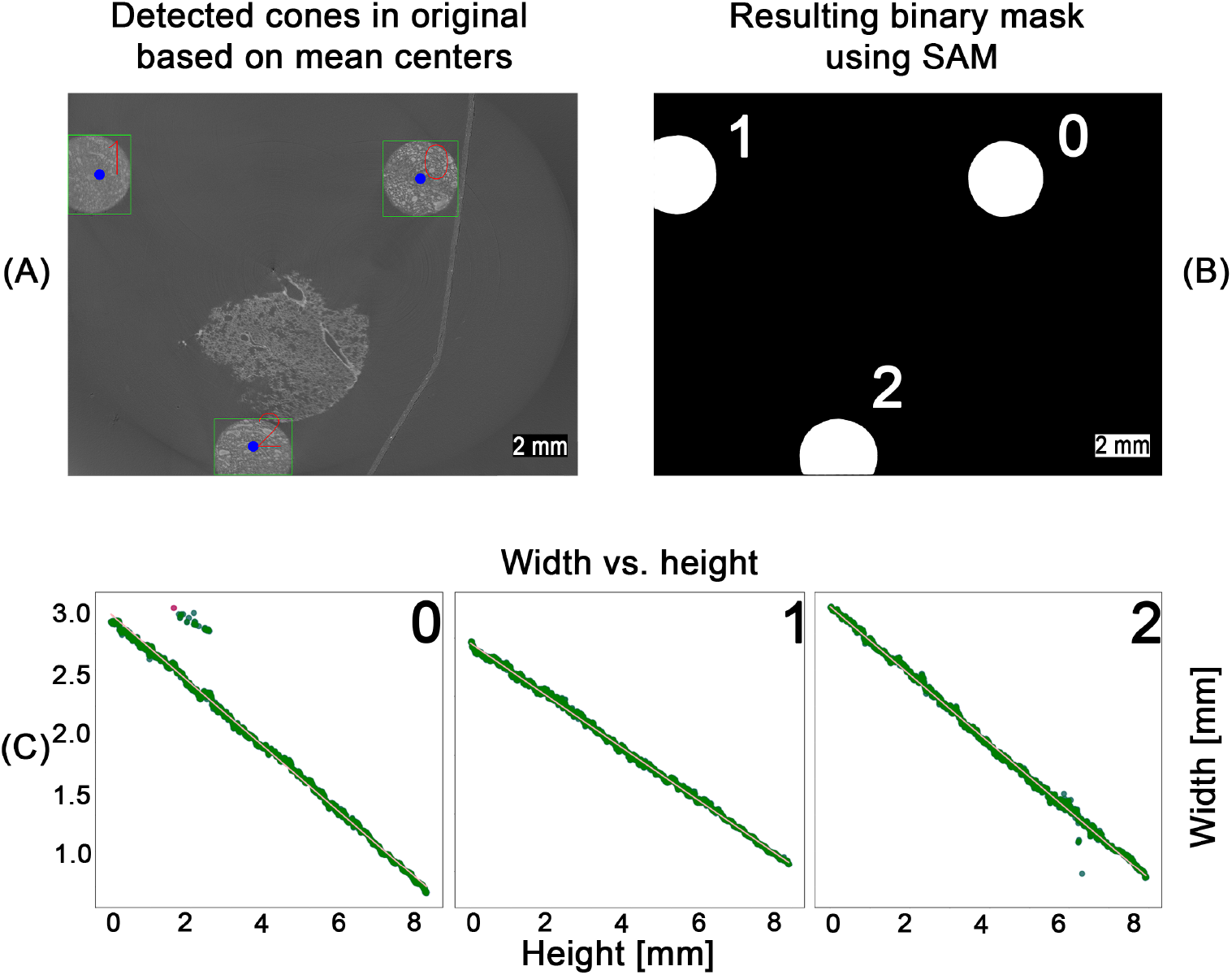
SAM-based detection workflow across the entire pipeline. Using the computed center points for each cone (blue dots), zero-shot detections from the SAM model were filtered (A). The resulting binary masks (B) were used to measure cone width at different heights. Width values were converted to millimeters and smoothed via linear regression (C), enabling mapping of the height–width relationship for each cone. Outliers (red dots), identified by their deviation from the regression line, were removed. The remaining inliers (green dots) were then used for a second linear regression to obtain the final line fit.

Based on the regression graphs for each cone, a computed width could be assigned to each layer of the z-stack, enabling direct comparison with the corresponding histological sections.

### Correlating PBI-microCT and histology

Following the algorithm described before [10], the histological image was processed using an image-processing pipeline to extract ellipse parameters. For this study, the method was extended with an automated matching step that assigned each histological ellipse to its corresponding cone based on approximate relative pixel coordinates. Using the measured major axes of the conic sections in the histology image, the corresponding layers in the PBI-microCT scan were identified via the regression graphs computed in the previous step (see Fig. 4). This yielded to the identification of the position of the individual conic section within the coordinates of the microCT based *in silico* volume. A cutting plane through the microCT data was defined using the three identified positions acting as a coarse initial guess for the subsequent image registration with the histological section. Image intensity values on this plane were sampled via trilinear interpolation, allowing for sub-voxel accurate resampling of the original image data. This approach enables flexible reorientation and precise extraction of oblique slices from volumetric datasets, thereby facilitating accurate image registration with histological sections. The extracted *in silico* plane was then rigidly registered to the histological section, as shown in Fig. 5.

**Fig. 5.**
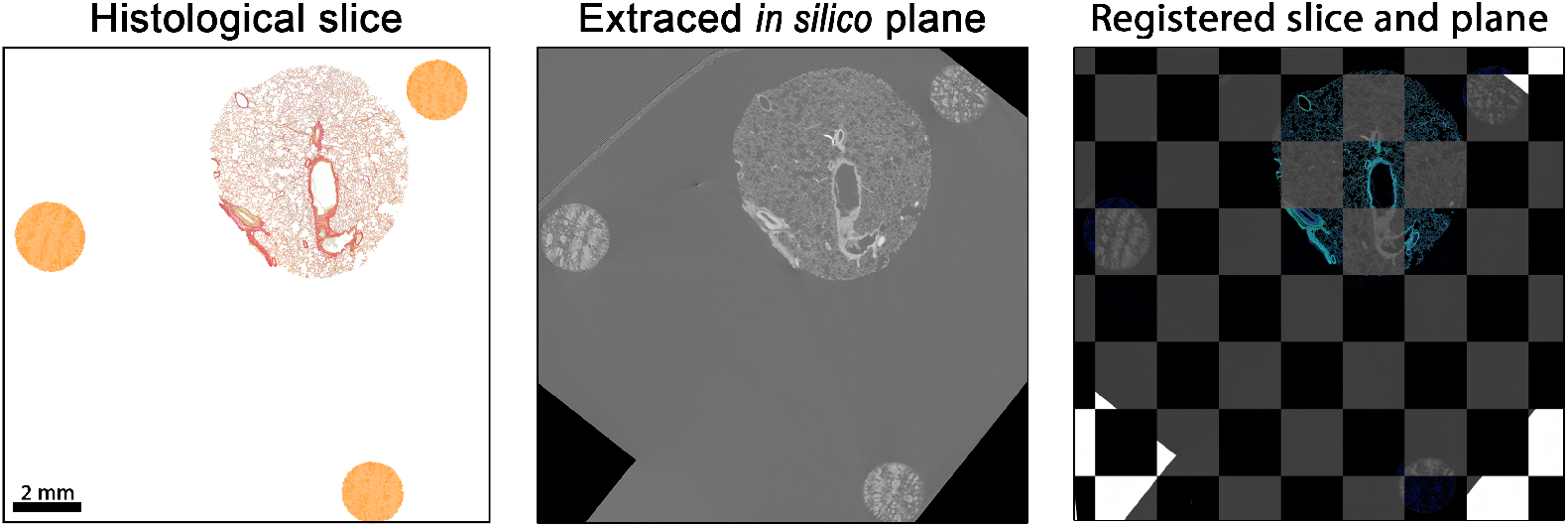
Matched and registered histological section and PBI-microCT plane. The histological slice was rigidly transformed to align with the extracted *in silico* plane by optimizing the mean square error, yielding an optimal spatial match. For better visual comparison, the colors of the histological section were digitally enhanced. Black pixels visible in the *in silico* plane result from rotation moving the plane partially outside the volume during the extraction process.

The resulting alignment provided a robust basis for precise matching. However, as parts of the cone structures extended beyond the field of view in the reconstructed PBI-microCT scan, subsequent registration steps focused primarily on the punch biopsy itself. Consequently, the final alignment was performed using only the tissue portion of the histological image. Using the methodology and registration tool introduced by Brettmacher et al. [31], the position of the extracted plane within the *in silico* volume served as the starting point for a 3D → 2D registration. The extracted image was then elastically aligned with the histological section to achieve precise correspondence.

To quantify the improvement in alignment, we employed the local normalized cross-correlation (LNCC) [40–43]. We chose a non-zero padding algorithm, as our analysis focused on the central regions of the image. Consequently, corner areas that would otherwise be influenced by padding effects were not fully included in the calculations. Furthermore, to prevent cancellation effects between positive and negative correlations, we modified the standard LNCC equation by computing the absolute value of each window-based normalized cross-correlation (NCC) score:

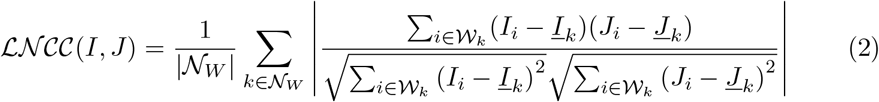

𝒩_*W*_ = set of all window centers,

𝒲_*k*_ = local window centered at pixel *k*,

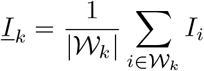 (mean intensity in the current window *W*_*k*_ in *I*),

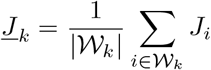 (mean intensity in the current window *W*_*k*_ in *J*),

*I*_*i*_, *J*_*i*_ = intensity values at voxel *i*.

This LNCC formulation is robust to local intensity and contrast variations. LNCC values were computed using a square window of size 45 *×* 45 pixels. For the analysis, we generated a heatmap in which each pixel represents the local NCC value calculated within its corresponding window. Due to the non-zero padding scheme, the resulting heatmap is smaller than the input image, with both height and width reduced by the selected window size. The mean of the heatmap corresponds to the overall LNCC value. In the resulting LNCC heatmaps (LNCC-HM), bright regions indicate high correlation, whereas dark regions indicate low correspondence. For improved visualization, the images were cropped to include only lung tissue. Fig. 6 shows LNCC-HMs for the histological section (A) aligned with: (B) a manually selected microCT plane, (C) a plane extracted using conic reference structures, and (D) a fully registered plane (rigid, affine, and elastic alignment). Panel (E) shows the final overlay between the fully registered microCT plane and the histological image.

**Fig. 6.**
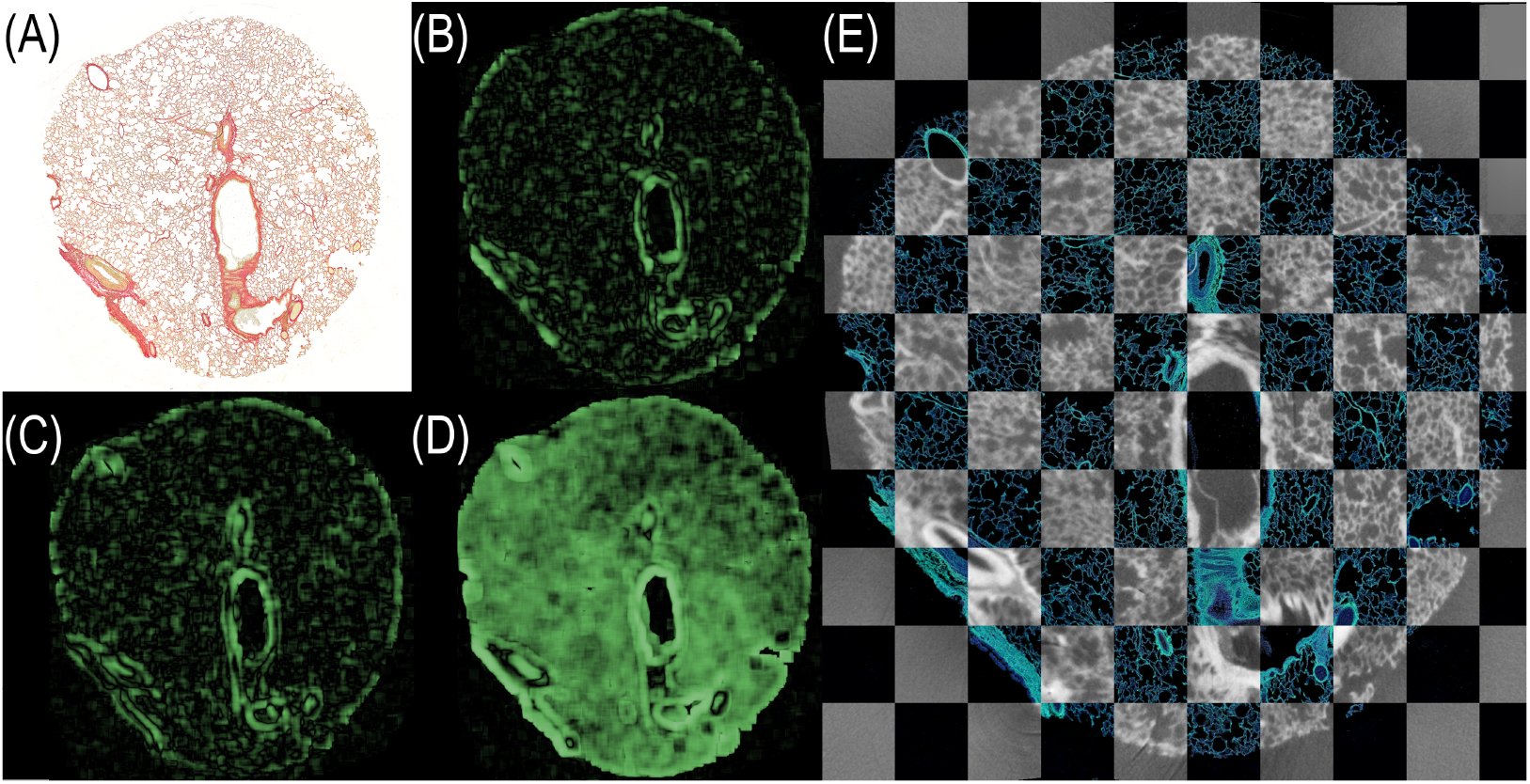
LNCC heatmaps visualizing the spatial correspondence between the histological section and various PBI-microCT planes. (A) Histological section. (B) Manually selected plane. (C) Plane extracted using conic reference structures. (D) Fully registered plane (rigid + affine + elastic). Green regions indicate strong local correlation; darker regions suggest misalignment. (E) Final overlay of fully registered microCT plane and histology depicted in a checkerboard pattern. The contrast of the histological section was enhanced for better visualization.

For our experiments we calculated the LNCC scores for five planes of the same volume. The results are listed in Table 1.

**Table 1.** Computed mean absolute LNCC values for three levels of registration precision. LNCC scores quantifying local spatial alignment between the histological section and microCT image at different registration stages for five planes.

| PBI-microCT image | Mean LNCC values | Max LNCC values |
| --- | --- | --- |
| Manual selection | $0.12 \pm 0.02$ | 0.14 |
| Extracted plane (conic reference) | $0.14 \pm 0.03$ | 0.19 |
| Fully registered plane | $0.29 \pm 0.09$ | 0.4 |

Both qualitative (visual) and quantitative (LNCC-based) evaluations demonstrate that the use of conic reference structures substantially improves plane extraction. Moreover, full image registration provides the most accurate alignment to the histological section.

## 4 Discussion

With the introduction of a novel resin and iodine-based contrast agent mixture, we achieved the generation of 3D-printed structures with tunable radiopacity. Building upon our earlier work [10], in which conical reference structures served as key tools for spatial correlation and multimodal fusion of microCT and histological imaging, we significantly improved both the manufacturing fidelity and image registration accuracy. Although the previously used cones were also 3D-printed, their coarse geometry limited registration precision. While sufficient for lower-resolution scans with voxel size of 80 *µ*m, they were inadequate for the higher-resolution PBI-microCT imaging used in this study, which achieves an isotropic resolution of 4 *µ*m. In this study we employed a different absorption-based microCT, which allows the reconstruction of a smaller field of view resulting in a voxel size of 18 *µ*m. However this approach results in visible noise preventing a precise overlay, as depicted in Fig. 2. We also evaluated the usage of the PBI-based Histomography blockscanner with a resolution of 10 *µ*m, to test the applicability of the conic markers in a setup independent of Synchrotron radiation. Across all scanners, our results demonstrate that the novel markers are suitable for high-resolution image fusion and can be readily segmented from the embedding medium due to their enhanced X-ray contrast. Our work also improved cone segmentation performance using the SAM2 [13], which provided robust results without requiring a training phase—offering a simple yet powerful alternative to other AI-based methods such as those presented by Chen et al. [2]. They utilized their CNN-based initialization [44] with the processed images 10 times smaller than used in our study. However, the LNCC achieved by the CNN method was smaller than the LNCC for the manual selection of the corresponding planes in the microCT data set. Despite the fact that the LNCC can not be directly compared to images of a different size, resolution and content, in our case for the fully registered plane, meaning automatic plane selection and elastic matching to the histological data, a technique comparable to the plane refinement used by Chen et al., the LNCC increased by more than a factor of 2 compared to manual selection.

Our method allows for the direct extraction of the corresponding *in silico* plane even if the orientation was tilted with respect to the slices of the microCT data stack, without the need for AI-based algorithms as shown in Nolte et al. [10]. The usage of SAM however increased the segmentation accuracy of the conic sections in the PBI-microCT slices from about 75% of the original approach to 100%. For the identification of the centers in those conic sections a YOLOv12 model [45] was implemented, which in the future could be trained to recognize conic sections in both microCT and histology. Beyond the registration aspect, our conic marker system was used for high-precision guided sectioning as shown in Nolte et al. [11]. Currently, the fusion process was limited by the comparable big size of the specimen, which necessitated performing multiple overlapping scans and introduced redundancy. In our experiments, some of the cones were only partially captured due to limited scan coverage. However, we did not observe any impairment in the ability to extract corresponding planes. Due to the slant angle of the cones, the difference of the major axes of the conic sections between each layer is limited. Thus, the accuracy of the correlation could be improved with increasing angles, which will be limited by the DLP printing resolution and would also increase the size of the cones. In the future, we plan to enhance the fusion process by incorporating morphological and colorimetric analysis from the histological sections, further demonstrating the value of successful multimodal correlation. Additionally, we envision the use of the resin-contrast agent mixture for producing radiopaque biomedical models from microCT data. These models, or digital twins, could be employed for educational, calibrational, or planning purposes [9, 46]. However, DLP currently requires parts to be printed separately and assembled manually, and does not support spatially graded radiopacity. To overcome these limitations, alternative resin-based printing technologies such as polyjet printing[47–49] could be explored.

For each matched image pair, we computed the LNCC score as well as a local NCC-based measure, which enabled the creation of a qualitative heatmap. This map highlights regions where the registration process performs well, as well as areas where performance could be improved. In future work, we aim to leverage these LNCC heatmaps (LNCC-HM) to guide targeted enhancements, ultimately improving the accuracy and quality of image overlays.

## 5 Conclusion

In conclusion, we introduced a novel 3D-printable resin-contrast agent mixture and demonstrated a streamlined, yet accurate approach for aligning PBI-microCT with histological imaging. This facilitates the generation of multimodal, spatially correlated datasets suitable for downstream machine learning applications without the need for manual labeling. Here previously unsupervised models could be finetuned. Moving forward, we aim to extend this methodology into a holistic tissue analysis framework and adapt the conical reference structures to other embedding media, such as paraffin.

## Author contributions

P.N. co-invented the resin–contrast agent mixture, designed the study, embedded and sectioned the specimens, and performed the majority of the software development. He also drafted the manuscript. C.J.A. co-invented the resin–contrast agent mixture, prepared and executed the 3D printing, and designed the reference structures. M.B. designed and performed the image registration and the image similarity analysis. M.R. scanned and reconstructed a subset of the specimens. M.S. provided the biological specimens. C.R. led the hardware development and secured funding. C.D. led the software development and microCT acquisition. All authors read, edited, and approved the final manuscript.

## Acknowledgments

The authors would like to acknowledge Ute Kant for her support in generating some of the slides and thank Rose Haarmann for her assistance in creating Fig. 1. Furthermore, the authors express their gratitude to Anfisa Eberle for her creative guidance.

## Data availability

The datasets used and/or analyzed during the current study are available from the corresponding author on reasonable request.

## Declarations

## Ethics declaration

This research project adheres to the highest ethical standards, and the principles of integrity and responsibility were rigorously upheld throughout the study. The nature of this investigation involved no human subjects, and no sensitive or confidential information was utilized. The animal experiments were performed and approved under project license AVD5020020209404 which was issued by the competent national authorities (Central Committee for Animal Experiments).

## Consent for publication

This manuscript hasn’t been published before and is not being considered for publication elsewhere. All the authors have contributed to the creation of this manuscript for important intellectual content and read and approved the final manuscript.

